# Towards intelligent living facades: On electrical activity of ordinary moss *Brachythecium rutabulum*

**DOI:** 10.1101/2024.04.28.591491

**Authors:** Andrew Adamatzky

## Abstract

Mosses display resilience and ecological importance, significantly shaping their environments. With their strong attachment to challenging substrates, mosses can serve as viable options for green living facades. In our initial steps towards developing sensing and computing living facades using moss, we analysed the endogenous electrical activity of mosses to establish foundational knowledge for future information processing devices. Employing macro-electrode recording techniques, we identified three patterns of electrical activity in ordinary moss: high-frequency oscillations at 1.2 Hz, medium-frequency oscillations at 2 · 10^−4^ Hz, and low-frequency oscillations at approximately 4 · 10^−4^. Additionally, we observed indications of coordinated electrical activity in moss cushions.

## 1. Introduction

Living facades, also known as green facades or vertical gardens, are architectural features where living vegetation is intentionally grown on the exterior surfaces of buildings [1, 2, 3, 4, 5, 6, 7, 8, 9, 10, 11]. These facades can range from simple climbing plants on trellises to complex systems with integrated irrigation and support structures. The purpose of living facades extends beyond mere aesthetics; they offer several environmental, economic, and social benefits, including biodiversity support, storm-water management, noise reduction, improved micro-climate, health and well-being. Living facades offer a multifaceted approach to sustainable urban design, integrating nature into the built environment to enhance aesthetics, mitigate environmental impact, and improve the quality of life for inhabitants and visitors alike. One approach to developing living facades involves synthesising bio-receptive concrete, which encourages living organisms to attach themselves to the concrete surface and successfully propagate [12, 13, 14, 15, 16, 17]. Another approach could involve sourcing living organisms with pre-existing strong attachment properties. Moss, is one of the potential candidates. Mosses, categorised under Bryophyta in the plant kingdom, are non-flowering plants distinguished by spore production, stems, and leaves. They lack true roots but wield significant ecological influence alongside their botanical relatives, liverworts and hornworts. Mosses demonstrate resilience and ecological significance, shaping their surroundings profoundly (Fig. 1a–c). Research was conducted on the rhizoids of various British moss species to explore their mechanical role in attaching to the substrate [18]. It has been demonstrated that the robust mechanical strength offered by the elongated rhizoids of mosses, along with the potential secretion of adhesive substances by these rhizoids, ensures effective anchoring of the shoots to challenging substrates. Excellent systematic study [19] demonstrated that mosses, with their high vegetative desiccation tolerance and minimal growing needs, can serve as an economical and easy-to-maintain substitute for green envelope systems, with suitable horizontal growth facilitated by capillary matting, cement, and lime plaster, although their vertical application is constrained by water distribution limitations.

**Figure 1:**
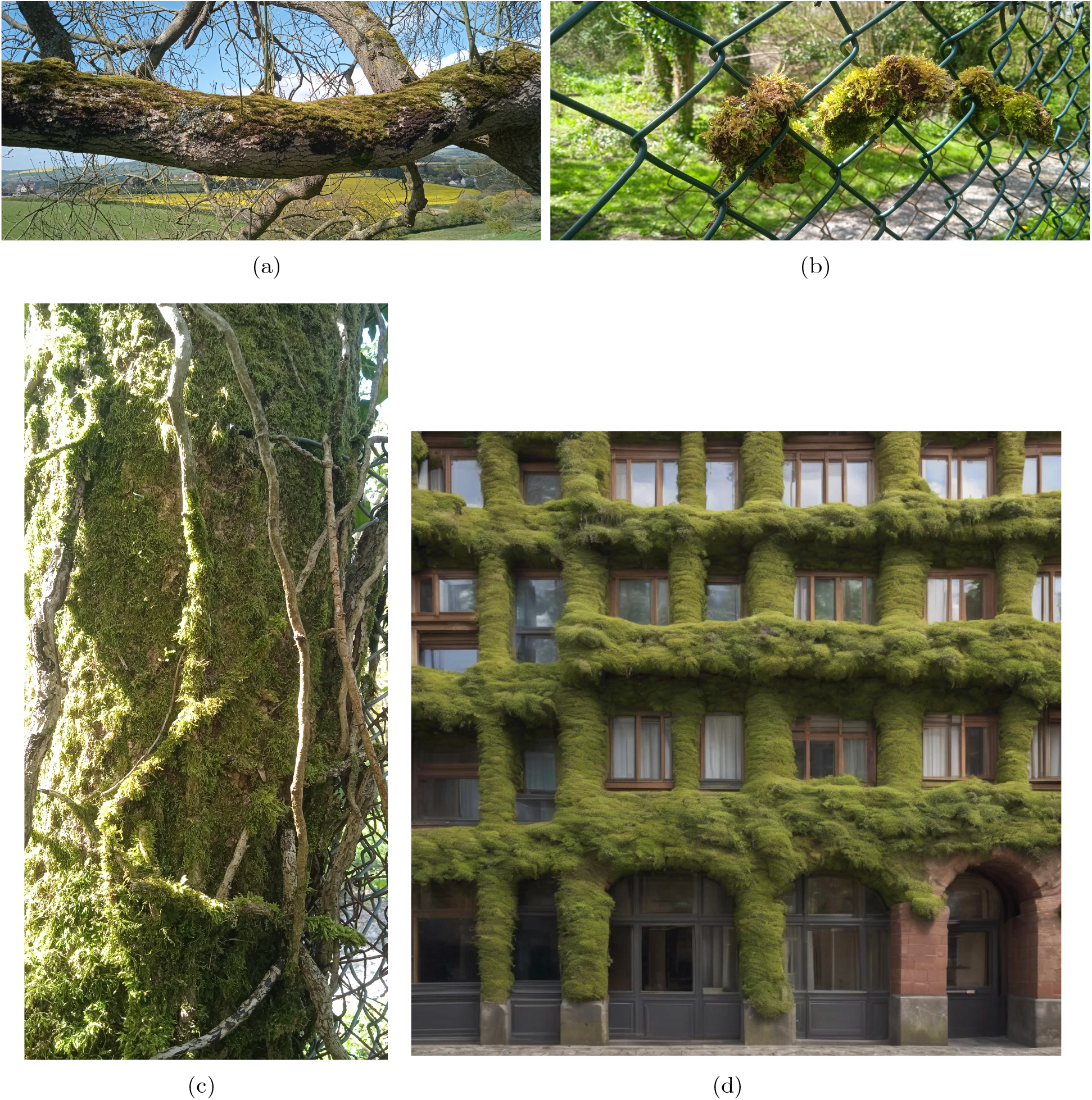
Moss growing on tree branches (a), wire fence (the moss isn’t merely hanging on the fence; it’s actually growing around the wires of the mesh) (b) and tree trunk (c). Photos are made in North Somerset, UK. (d) Artistic impression of the moss-based responsive living facade.

Future living facades constructed using moss (as depicted in Fig. 1d) are poised to function not only as responsive passive screens but also as adaptive learning facades, potentially serving as expansive sensory networks and biocomputers.

An interface utilising living sensing and computing facades can be implemented through optical, chemical, and electrical methods. Assuming that the nature of both the input and output interfaces should be consistent, electrical communication emerges as the most advantageous option. As an initial step towards exploring the information processing properties of moss facades, we have decided to classify patterns of electrical activity in moss.

Plants are renowned for their salient intelligence [20, 21, 22, 23, 24], capabilities to implement distributed information processing [25, 26, 27, 28], showing indicators of advanced perception, cognition and adaptive behaviour [29, 30, 31, 32], anticipatory responses [33] and swarm intelligence [34]. Plants employ impulses of electrical activity to coordinate actions of their bodies and long-distance communication [35, 36, 37]. The bursts of impulses could be either endogenous [38], e.g. related to motor activities [39, 40, 41, 42] or in a response to external stimulation, e.g. temperature [43], osmotic environment [44], mechanical stimulation [45, 46]. Electrical signals could propagate between any types of cells in a plant tissue, there are indications however of higher conductivity of the vascular system which might act as a network of pathways for travelling electrical impulses [47, 48, 49, 50].

The electrical activity of moss is scarcely studied; however, the few papers published on the topic show very interesting and encouraging results. Light induced action-potential like membrane potential depolarisation has been shown in [51, 52]. The research presented in [53] demonstrates that moss responds to thermal and chemical stimulation by generating electrical potential spikes with distinctive shapes. Long distance signalling by changed in the membrane potential in response to glutamate stimulation has been demonstrated in [54]. Most of the experiments mentioned have been conducted using micro-electrode measurements. We have decided to adopt macro-measuring techniques as they are the most feasible and economical option in the context of real-life interfacing with moss facades. The technique is presented in Sect. 2, results obtained are analysed in Sect. 3 and discussed in Sect. 4.

## 2. Methods

The electrical activity of the kombucha was recorded using pairs of iridium-coated stainless steel sub-dermal needle electrodes (Spes Medica S.r.l., Italy), connected to twisted cables and an ADC-24 high-resolution data logger from Pico Technology (UK) equipped with a 24-bit analog-to-digital converter (see Fig.2a). Data were recorded at a rate of one sample per second, with the logger capturing as many measurements as possible (typically up to 600 per second) and saving the average value. The acquisition voltage range was set to 156 mV with an offset accuracy of 9 μV at 1Hz to maintain a gain error of 0.1%. Each electrode pair, treated as independent channels, reported the difference in electrical potential between the electrodes. The distance between electrodes ranged from 1 to 2 cm. Data were collected from eight pairs of differential electrodes during uninterrupted recording for 178 h (just over seven days), resulting in over 5 10^6^ measurements analysed (see Fig.2c). The moss container was kept in a transparent closed container with high humidity, in a room with constant ambient lighting of approximately 5 lux.

**Figure 2:**
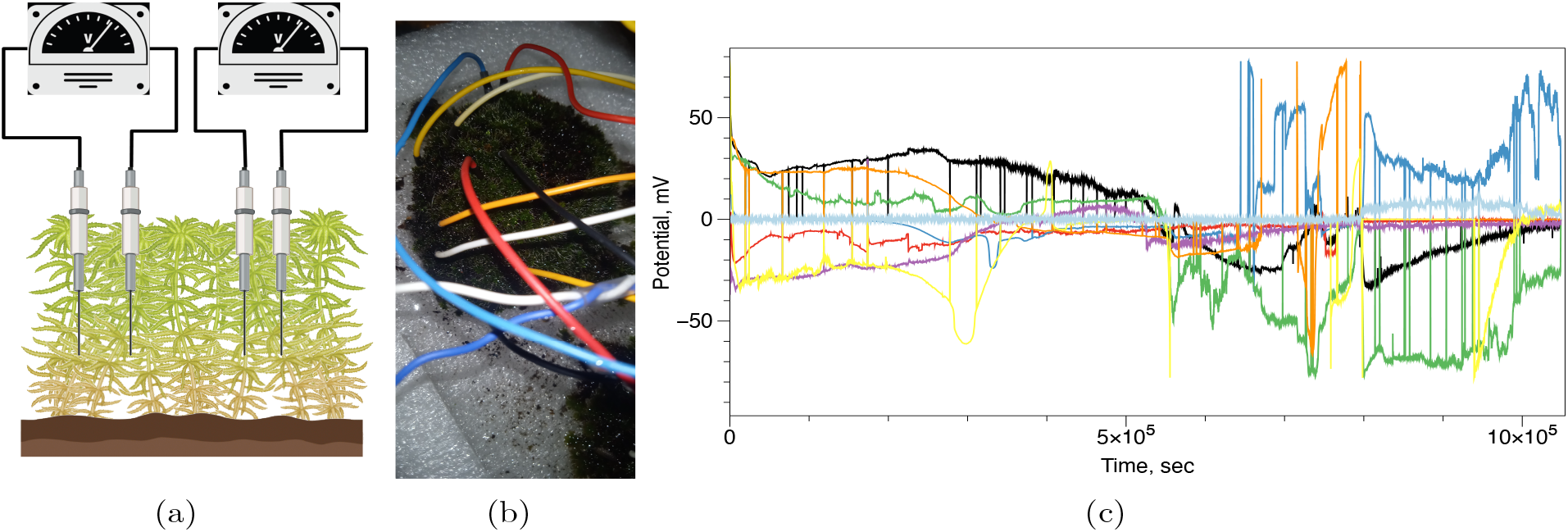
Experimental setup: (a) Scheme depicting moss *B. rutabulum* on a wet substrate with two pairs of visible differential electrodes. (b) A photo taken from above showing the moss and electrodes’ cables. (c) Overview of the electrical activity of *B. rutabulum*. Recordings from each pair of differential electrodes are represented by a unique color.

## 3. Results

We found that the moss *B. rutabulum* exhibits a diverse range of extracellular potential spiking activity, categorized into three distinct patterns: high-frequency spiking (Fig.3a), medium-frequency spiking (Fig.3b), and low-frequency spiking. The qualitative statistical characteristics of these patterns are detailed in Tab. 1a.

**Figure 3:**
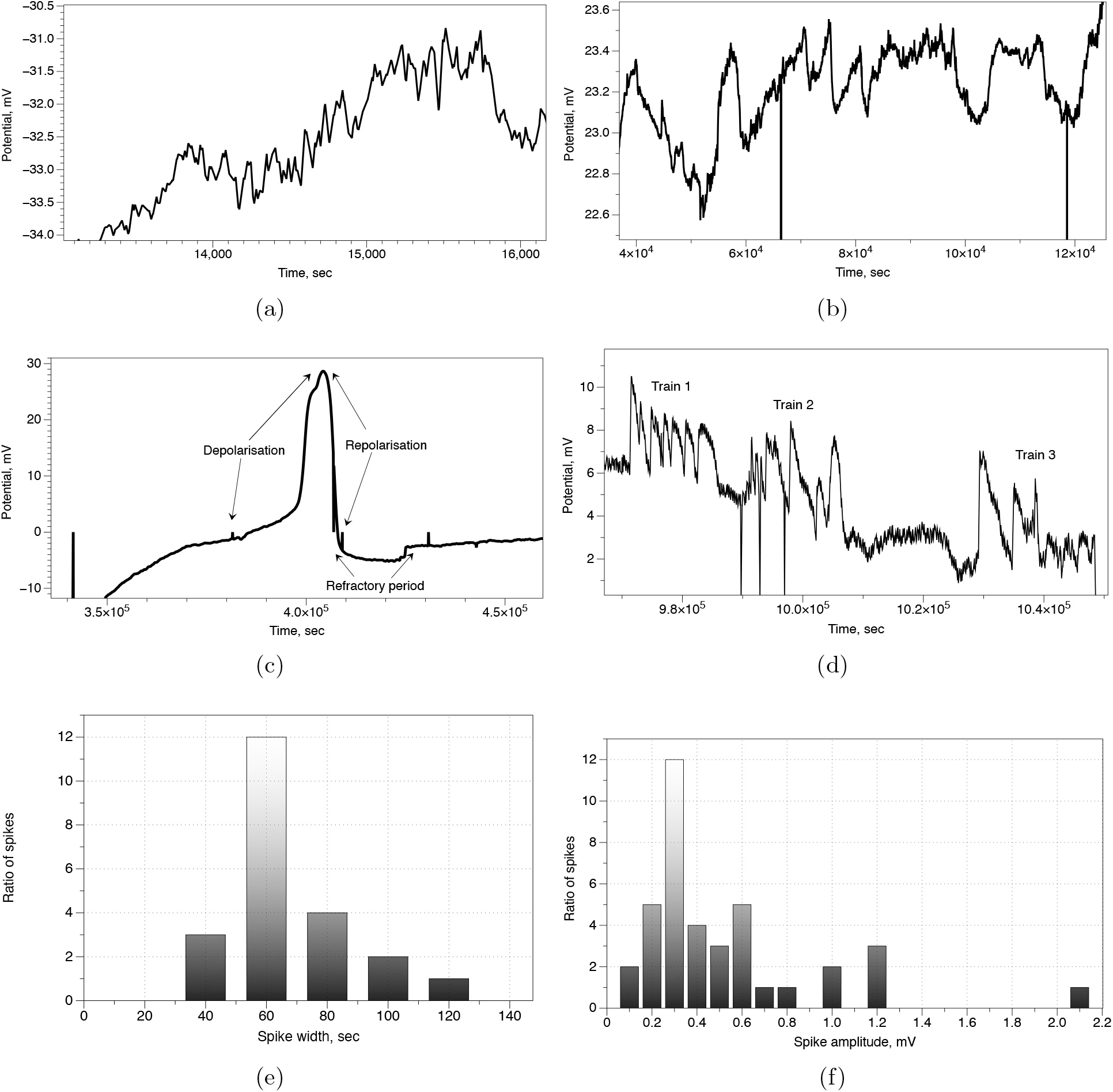
Examples of electrical spiking activity in *B. rutabulum*. (a) High-frequency spikes. (b) Low-frequency spikes. (c) Large action-potential like spike. (d) Distribution of spike amplitudes. Bin size is 0.1. (e) Distribution of width of fast spike. Bin size is 20.

**Table 1:**
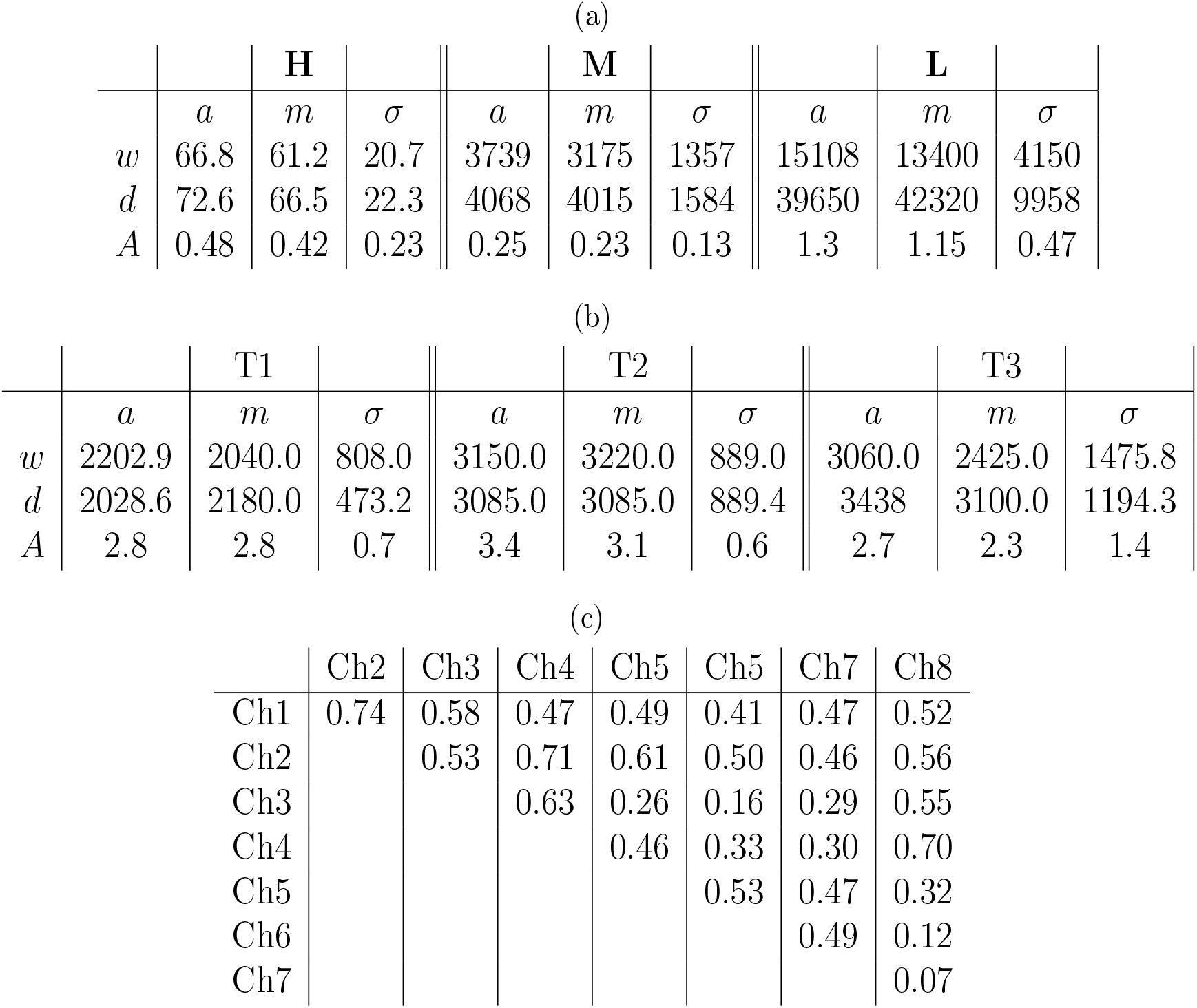
Characteristics of electrical spiking patterns recorded in *B. rutabulum* (a) **H** are high-frequency spikes, **M** medium-frequency spikes, and **L** are low frequency spikes. (b) Characteristics of spike trains, shown in Fig. 3d. First, second and third trains are presented as T1, T2 and T3. *w* is a spike width in sec, *d* is a distance between consecutive spikes in sec, *A* is a spike amplitude in mV, *a* is an average value, *m* is a median value and *σ* is a standard deviation. (c) Absolute values of correlation coefficients between channels.

|  | **H** |  |  |  | **M** |  |  |  | **L** |  |  |
| --- | --- | --- | --- | --- | --- | --- | --- | --- | --- | --- | --- |
| | *a* | *m* | $\sigma$ | | *a* | *m* | $\sigma$ | | *a* | *m* | $\sigma$ |
| *w* | 66.8 | 61.2 | 20.7 |  | 3739 | 3175 | 1357 |  | 15108 | 13400 | 4150 |
| *d* | 72.6 | 66.5 | 22.3 |  | 4068 | 4015 | 1584 |  | 39650 | 42320 | 9958 |
| *A* | 0.48 | 0.42 | 0.23 |  | 0.25 | 0.23 | 0.13 |  | 1.3 | 1.15 | 0.47 |

(b)
|  | **T1** |  |  |  | **T2** |  |  |  | **T3** |  |  |
| --- | --- | --- | --- | --- | --- | --- | --- | --- | --- | --- | --- |
| | *a* | *m* | $\sigma$ | | *a* | *m* | $\sigma$ | | *a* | *m* | $\sigma$ |
| *w* | 2202.9 | 2040.0 | 808.0 |  | 3150.0 | 3220.0 | 889.0 |  | 3060.0 | 2425.0 | 1475.8 |
| *d* | 2028.6 | 2180.0 | 473.2 |  | 3085.0 | 3085.0 | 889.4 |  | 3438 | 3100.0 | 1194.3 |
| *A* | 2.8 | 2.8 | 0.7 |  | 3.4 | 3.1 | 0.6 |  | 2.7 | 2.3 | 1.4 |

Table 1: Characteristics of electrical spiking patterns recorded in *B.
|  | Ch2 | Ch3 | Ch4 | Ch5 | Ch5 | Ch7 | Ch8 |
| --- | --- | --- | --- | --- | --- | --- | --- |
| Ch1 | 0.74 | 0.58 | 0.47 | 0.49 | 0.41 | 0.47 | 0.52 |
| Ch2 |  | 0.53 | 0.71 | 0.61 | 0.50 | 0.46 | 0.56 |
| Ch3 |  |  | 0.63 | 0.26 | 0.16 | 0.29 | 0.55 |
| Ch4 |  |  |  | 0.46 | 0.33 | 0.30 | 0.70 |
| Ch5 |  |  |  |  | 0.53 | 0.47 | 0.32 |
| Ch6 |  |  |  |  |  | 0.49 | 0.12 |
| Ch7 |  |  |  |  |  |  | 0.07 |

High-frequency spiking patterns consist of spikes approximately 67 seconds wide (Fig. 3e), occurring at a frequency of around 1.2 Hz, with amplitudes of approximately 0.4 mV. These spikes typically do not occur in trains.

Medium-frequency spikes have a typical duration of about 1 hour, with an oscillation frequency around 2 10^−4^ Hz. They exhibit smaller amplitudes (Fig. 3f), averaging about 0.25 mV.

Low-frequency patterns of electrical activity have an average spike duration of approximately 4.2 hours and an oscillation frequency around 7 10^−5^.

In addition to these described classes, we also observed spikes resembling high-amplitude action potentials and neuron-like spike trains.

A high-amplitude spike with action potential characteristics is depicted in Fig. 3c. The depolarisation phase lasts 6.2 hours, during which the potential increases by 30 mV. The repolarisation phase lasts 1.3 hours, during which the potential decreases by 31.2 mV. The refractory period is 2.8 hours, with a potential drop of 2.5 mV. These large spikes occur with an average interval of 18 hours.

Three trains of electrical potential spikes are depicted in Fig.3c: the first train consists of seven spikes, the second train has five spikes, and the third train comprises six spikes. The quantitative characteristics of these trains are presented in Tab.1b. The duration of a spike in the trains varies from 0.6h to 0.8h and the frequencies of oscillations range from approximately 3 · 10^−4^ to 5 · 10^−4^ Hz.

Correlation coefficients between recording channels are shown in Tab. 1c, an exemplar scattering plot of two neighbouring channels in Fig. 4a and distribution of the absolute values is shown in Fig. 4b. Over half of the coefficients are above 0.5. The extent of the correlation being observed indicates a substantial level of synchronisation or coordination in the electrical activity across different parts or aspects of the moss being studied.

**Figure 4:**
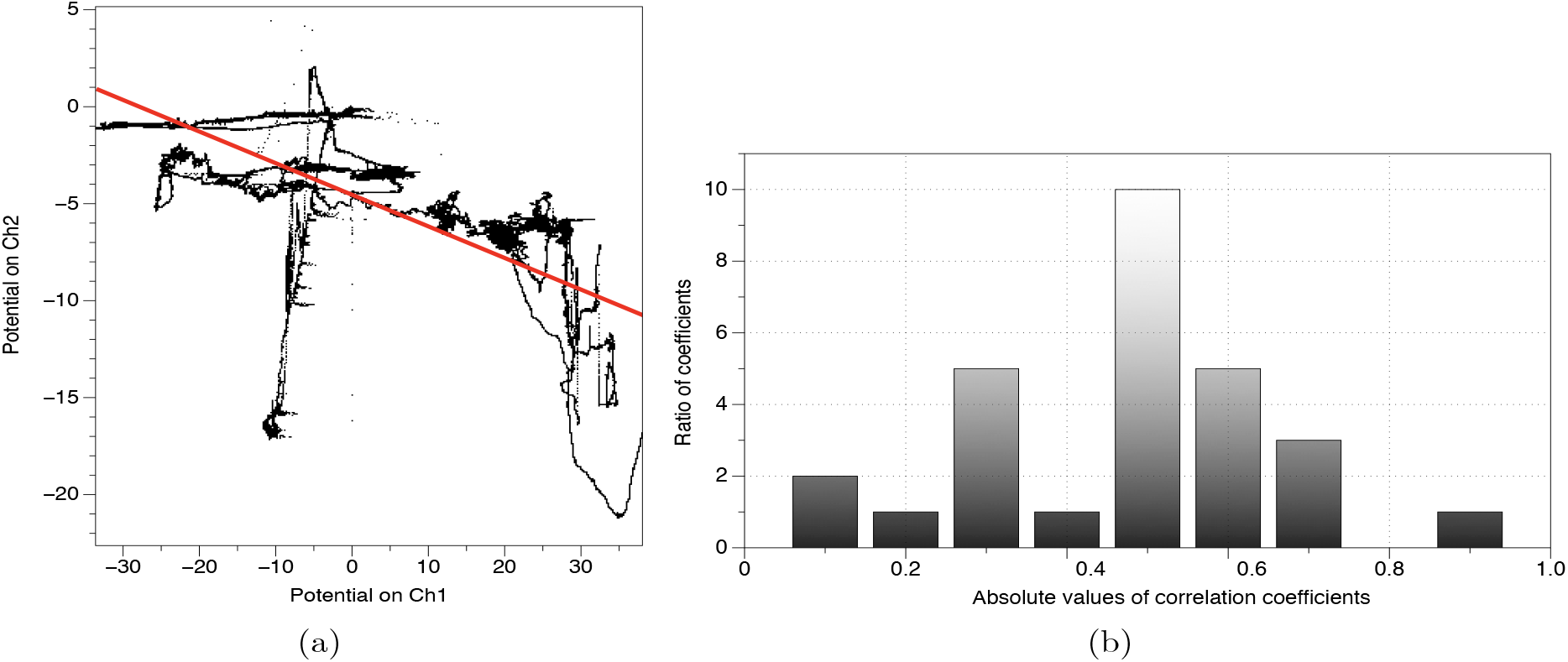
Correlations between recording channels. (a) Scattering plot of two neighbouring channel with linear approximation, show by red line, *Ch*2 = (− 4.5401) + (− 0.16296) * *Ch*1. (b) Distribution of absolute values of correlation, between recording channels, coefficients.

## 4. Discussion

In experimental laboratory setup we uncovered three families of electrical activity pattern of ordinary moss. These are high-frequency oscillations of 1.2 Hz, medium-frequency oscillations of 2· 10^−4^ Hz and low-frequency oscillations of c. 4 · 10^−4^. Based on the illustrations of action-potential like spikes in micro-electrode recordings [51, 52, 53, 54] we can speculate that high-frequency patterns of periodic electrical activity originate from integrated oscillations of individual plants, possibly synchronised or coupled via rhizoids. One potential explanation could involve the activation of ion channels in response to rapid changes in environmental conditions, such as changes in light intensity or temperature fluctuations. However, the experimental space remained in stable illumination and temperature conditions. High-frequency oscillations might also be linked to rapid cellular processes, such as action potentials or membrane depolarisation events, occurring within the moss cells.

The medium-frequency oscillations could be associated with slower regulatory mechanisms within the moss, such as ion channel gating kinetics or regulatory feedback loops.They might also reflect oscillatory behaviours arising from interactions between different cellular compartments or signalling pathways. The low-frequency oscillations might be driven by slower physiological processes, such as nutrient uptake and transport, metabolic cycles, or circadian rhythms inherent to the moss. They could also be influenced by external factors with longer time scales, such as diurnal variations in environmental conditions or the moss’s response to seasonal changes.

Possible explanations and mechanisms for the medium- and low-frequency spiking patterns could involve various factors such as the intrinsic properties of moss cells, interactions with environmental stimuli like light or moisture, regulatory mechanisms within the moss’s signalling pathways, and potential interactions with microorganisms present in the moss ecosystem. Further investigation into these factors and their interplay could shed light on the underlying mechanisms driving the observed electrical activity patterns.

